# Study on the phenotypic diversity and comprehensive evaluation analysis of 43 ornamental peonies of Sect. *Paeonia*

**DOI:** 10.1101/2024.08.06.606934

**Authors:** Hui-yan Cao, Shi-yi Xu, Mei-qi Liu, Shan Jiang, Leng-leng Ma, Jian-hao Wu, Xiao-Zhuang Zhang, Ling-yang Kong, Wei-chao Ren, Zhi-yang Liu, Xi Chen, Wei Ma, Xiu-bo Liu

**Affiliations:** School of Pharmacy, Heilongjiang University of Chinese Medicine, Harbin 150040, China; Experimental Training Center, Heilongjiang University of Chinese Medicine, Harbin 150040, China; Harbin Academy of Agricultural Sciences, Harbin 150028, China; Jiamusi College, Heilongjiang University of Chinese Medicine, Jiamusi 154007, China

## Abstract

The peony of Sect. *Paeonia* was a perennial herbaceous plant with numerous ornamental varieties and riched diversity in flower color and shape. It has ornamental, edible, and medicinal value and a long history of cultivation in China. The study of phenotypic diversity of plants is an important foundation for plants of Sect. *Paeonia* breeding. This study conducted phenotypic diversity analysis, principal component analysis, and cluster analysis on 43 varieties of Sect. *Paeonia* germplasm resources. Phenotypic traits included 30 qualitative traits and 7 quantitative traits. Through genetic diversity analysis, principal component analysis, comprehensive evaluation, and cluster analysis, we ultimately concluded that plant samples had relatively rich genetic phenotype traits. In principal component analysis, the first 12 principal components have covered the vast majority of information for phenotypic traits. The comprehensive evaluation results of phenotypic traits indicate that the F values of each variety in the germplasm sample were all positive number. The degree of stamen petals played a key role in determining the phenotypic diversity of plants, and the shape of the cotyledons and leaflets may determine the plant’s stress resistance performance, which provides a reference for breeding new varieties of peonies of Sect. *Paeonia*.

## Introduction

Sect. *Paeonia* is a perennial herbaceous flowera species belonging to the genus *Paeonia.* in the *Paeoniaceae* family [1,2]. The plants of genus *Paeonia* are divided into three groups, namely the Sect. *Moutan* [3], the Sect. *Oanopia*, and the Sect. *Paeonia* [4]. Plants of Sect. *Paeonia* are colorful and diverse [5], in addition, the apparent diversity of plants is also very rich [6]. The plants of genus *Paeonia* have a long cultivation history and abundant planting resources in China. In recent years, the plants of genus *Paeonia* have received widespread attention in the cut flower market, landscaping applications, and the development of nutritional and health products [7–11] and it has high economic and ornamental value.

The combined effect of genetic material and growth environment regulates the phenotypic traits of plants, and plants gradually form stable external manifestations through long-term natural selection [12,13]. Different varieties of ornamental plants of Sect. *Paeonia* have different physiological and biological characteristics, and its growth status show significant differences in different regions^14^. Phenotypic traits are important indicators of adaptation to the environment, selection, survival, and evolution [15,16]. The study of phenotypic traits can provide an overall understanding of the genetic diversity of research subjects, which is a reflection of individual genetic diversity and environmental differences within plant species [17–20].

The genetic basis of plants of Sect. *Paeonia* are relatively complex, and there is a significant or extremely significant correlation between each trait [21,22]. The differences in plant introduction and domestication between different varieties, especially in terms of morphological and physiological characteristics, worth more attention from researchers. The introduction and domestication [23,24] of plants are the main means to enrich local plant resources, as well as an important foundation for genetic improvement and new variety cultivation. Establishing an accurate and effective classification and identification system, as well as clarifying the genetic relationships between varieties [25,26] is crucial for hybrid breeding and plant resource development and utilization during introduction and domestication, and is crucial for product quality control. The parents of artificially cultivated plants of Sect. *Paeonia* have become more complex through continuous domestication and cultivation, and the genetic diversity phenotypes of offspring are more abundant [27–29]. Of course, different ecological environments can also lead to rich genetic variations. Analyzing the diversity and investigating the survival status of the plants of Sect. *Paeonia* is beneficial for resource conservation and the development of new varieties.

## Materials and methods

### Materials

The test materials were provided by Germplasm Resource Nursery of Harbin Academy of Agricultural Sciences, Heilongjiang Province (126o55′ E, 45o8′ N). The plant resources of the Sect. *Paeoni*a we have chosen were all perennial and well growing varieties.The altitude range of the nursery was between 115 and 120 meters. The annual average temperature was 4.2 ℃, with a maximum temperature of 36.7 ℃ and a minimum temperature of -37.7 ℃. The annual fros tfree period can last for 140-160 days, the maximum snow depth can reach 41cm. The relative humidity of the air was about 66%, the average precipitation is about 524.5mm, and the annual average evaporation was about 1586.8mm. The planting spacing of the plant nursery should be controlled at 50×50. The plant samples selected for this experiment were 6-year-old peony varieties, totaling 43 varieties. Phenotypic trait parameters were collected through observation and measurement. 10 plants were randomly selected from the germplasm samples of each variety as research subjects, and the phenotypic trait data of each plant was collected multiple times to retain 3 parallel results.

### Methods

A total of 37 phenotypic traits were investigated, divided into two categories: qualitative traits and quantitative traits. Phenotypic traits included growth habit, flower posture, crown diameter, petiole color, plant height, and number of branches. The direct observation method was used for the qualitative traits of the sample, and the specific measurement method was used for the quantitative traits. The observation standards were based on the Agricultural Industry Standard of the People’s Republic of China (NY/T2225-2012—Guidelines for Specificity, Consistency and Stability Testing of New Plant Varieties. All color related traits in this study were based on the colorimetric card published by the Royal Horticultural Association of the United Kingdom as the reference standard. The tools required for collecting the observation data mainly included the colorimetric card, vernier caliper and tape measure.

### Data statistics and analysis

Multivariate statistical methods are used to evaluate the degree of genetic diversity performance of different varieties of peony. Excel 2016 was used to calculate the genetic diversity index of qualitative traits and process relevant experimental data, analyzed the phenotypic diversity index, coefficient of variation, and the average coefficient of variation of phenotypic characters of the samples.Assigning and grading traits based on the mean (X) and standard deviation(δ), the number of grades depends on the complexity or numerical range of germplasm resource traits.The difference in property weight between each level was 0.5 δ. Calculate the Shannon Wiener diversity index (*H* ’) using the formula:

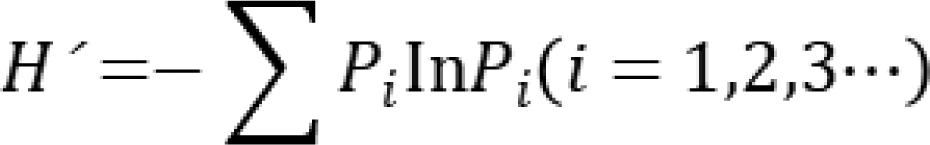

*P_i_* was the percentage of materials at the *i*-th level of a certain trait to the total number of materials. Principal component analysis (PCA) and cluster analysis were performed on the characteristics of the test materials using SPSS 19.0. In the PCA, principal components were extracted based on characteristic value >1. The standardized values of traits were input into each principal component, and the scores of the principal components were calculated. The property weight was determined based on the contribution rate of the principal components. Fuzzy membership functions were used to normalize each principal component, and the comprehensive evaluation values of all variety samples were obtained.

Cluster analysis was conducted using phenotypic traits and peony varieties as variables. ***SPSS 19.0*** software was used as the analysis tool, the range of the variable analysiswas set to a minimum of 2 and a maximum of 5. Cluster analysis used inter group linkage method based on *Euclidean distance*.

## Results

The names and codes of different plant varieties selected for research of Sect. *Paeonia* were listed in Table 1. The morphological images of all varieties were showed in Figure 1.

**Table 1.**
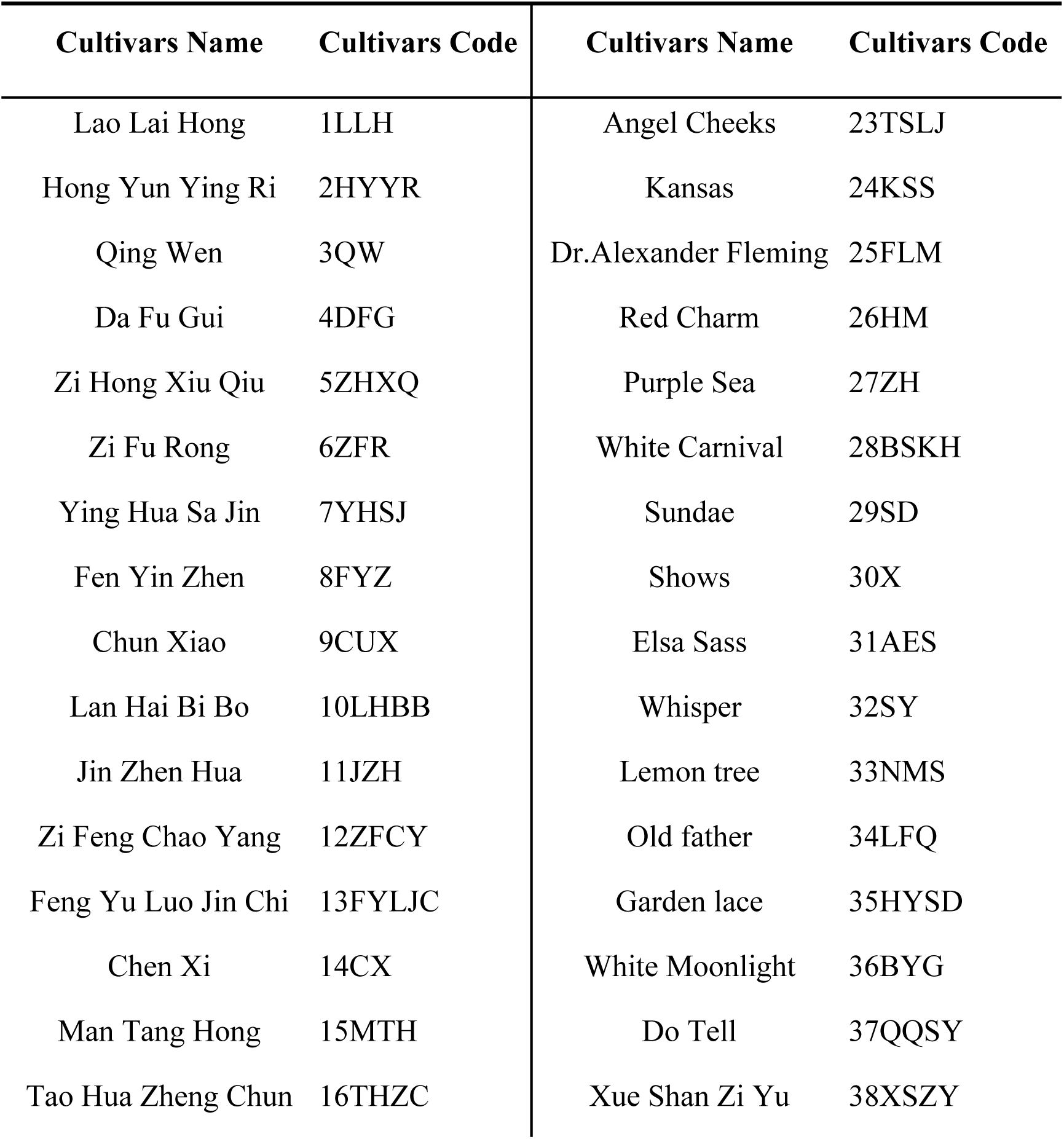
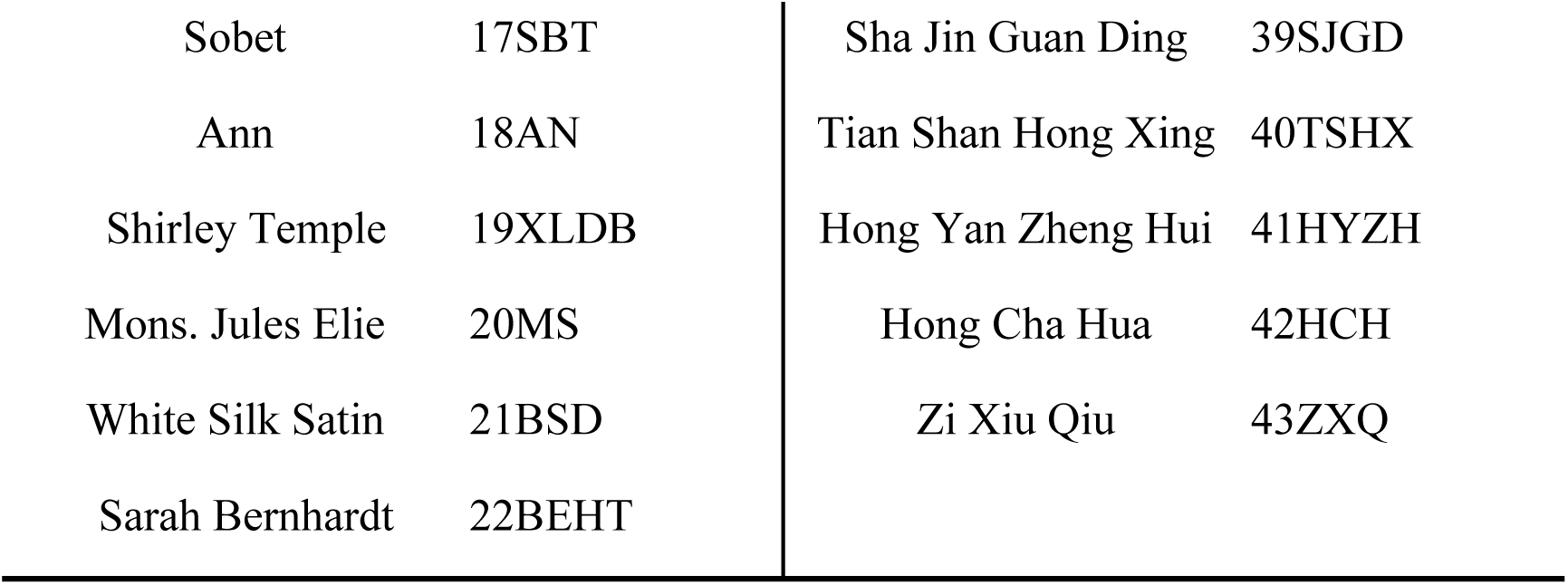
Names and codes of plant germplasm resources of Sect. *Paeonia*.

**Figure 1.**
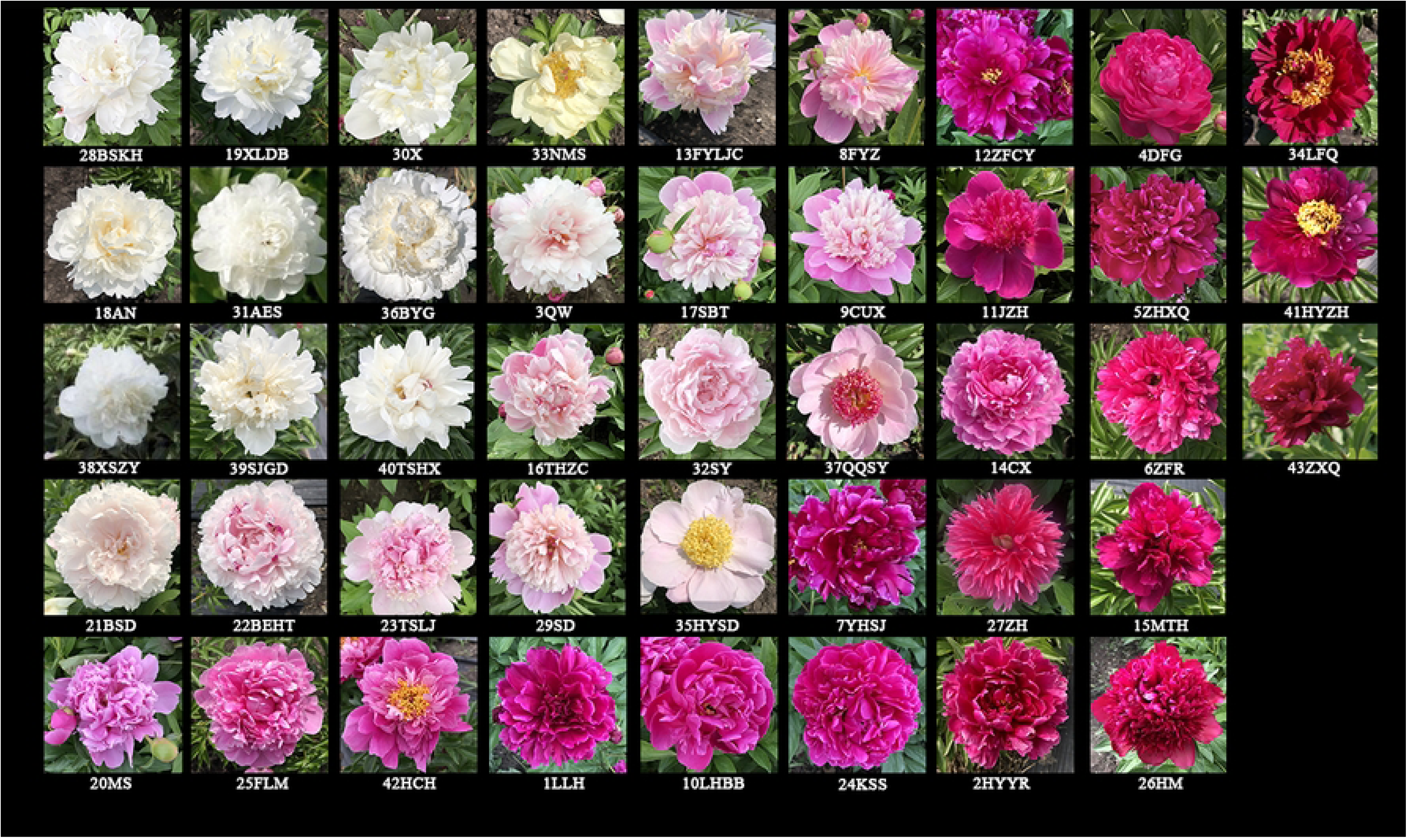
The morphological images of 43 varieties of peonies of Sect. *Paeonia*.

### Morphology parameters collection and phenotypic diversity analysis

The descriptive statistical analysis results of 30 qualitative traits of 34 cultivars were shown in Table 2 and Table 3, The phenotypic diversity index of the qualitative traits ranged from 0.018 to 0.500 (average=0.143). The compound leaves type, compound leaves petiole color, flower bud colors, flower shape, flower color, degree of petalization of stamens, stamen petal shape and stamen petal color revealed a phenotypic diversity index of >0.2. The compound leaves type had the highest *H’* (0.500), this indicates that there was a rich variation in the types of compound leaves among different varieties. The compound leaves type was mainly represented by the medium round shape, with a distribution frequency of 34.88%. Among them, the *H’* of flower color and degree of petalization of stamens were relatively high, the values are 0.471 and 0.392 respectively. Although there were many manifestations of the degree of petalization of stamens, most of them exhibit characteristics of petalization. Such as he distribution frequency of the epigenetic features of petalization of stamens was 41.86%, this indicated that in most varieties of peony plants, they choose to abandon their stamens and produce complex double petal flowers through petalization. However, this feature was not very strong in the process of pistil petalization, and the distribution frequency of pistil without petal variation among 43 varieties was as high as 76.74%. These qualitative traits with high *H’* were mainly characterized by plant color and shape, indicating that plant color and shape were the main factors contribut to the qualitative traits among different varieties. The *H’* of pistil petal color is the lowest (0.018), the *H’* of degree of pistil petalization was 0.043, and more than 75% of the samples did not have pistil petalization, which indicated that pistil petalization was hidden during the hybridization process.

**Table 2.** Analysis of diversity of qualitative characters of peonies of Sect. *Paeonia*.

| Traits | Phenotypic<br>Diversity Index<br>$H'$ | Coefficient of<br>Variation/% |
| --- | --- | --- |
| Growth habit | 0.064 | 39.691 |
| Plant height | 0.074 | 32.405 |
| Number of flowers | 0.063 | 30.241 |
| The dividing ability of the scale buds | 0.078 | 42.672 |
| Compound leaves type | 0.5 | 48.966 |
| Angle of compound leaf petiole and flower<br>branch | 0.102 | 30.071 |
| Compound leaves upper surface color | 0.149 | 32.027 |
| Compound leaves petiole color | 0.201 | 28.375 |
| Degree of inward curling of leaflet | 0.098 | 25.267 |
| Waviness degree of leaflet margin | 0.054 | 32.125 |
| Leaflet apex shape | 0.111 | 61.688 |
| Basic association of parietal leaflet | 0.022 | 24.096 |
| flower bud blooms | 0.023 | 26.889 |
| flower bud colors | 0.269 | 43.173 |
| Whether there is lateral bud | 0.045 | 35.454 |
| Flower branch diameter | 0.132 | 29.17 |
| Flower branch length | 0.082 | 31.104 |
| Flower shape | 0.284 | 48.352 |
| Flower posture | 0.091 | 35.243 |
| Pedicel color | 0.048 | 56.547 |
| Flower color | 0.471 | 51.389 |
| Whether there is sepal petalization | 0.08 | 26.372 |
| Outer petal shape | 0.09 | 47.482 |
| Degree of petalization of stamens | 0.392 | 38.934 |
| Stamen petal shape | 0.271 | 34.257 |
| Stamen petal color | 0.299 | 62.292 |
| Degree of pistil petalization | 0.043 | 84.014 |
| Pistil petal color | 0.018 | 71.58 |
| Color of stamen filaments | 0.064 | 31.565 |
| Color of pistil stigma | 0.063 | 76.538 |

**Table 3.**
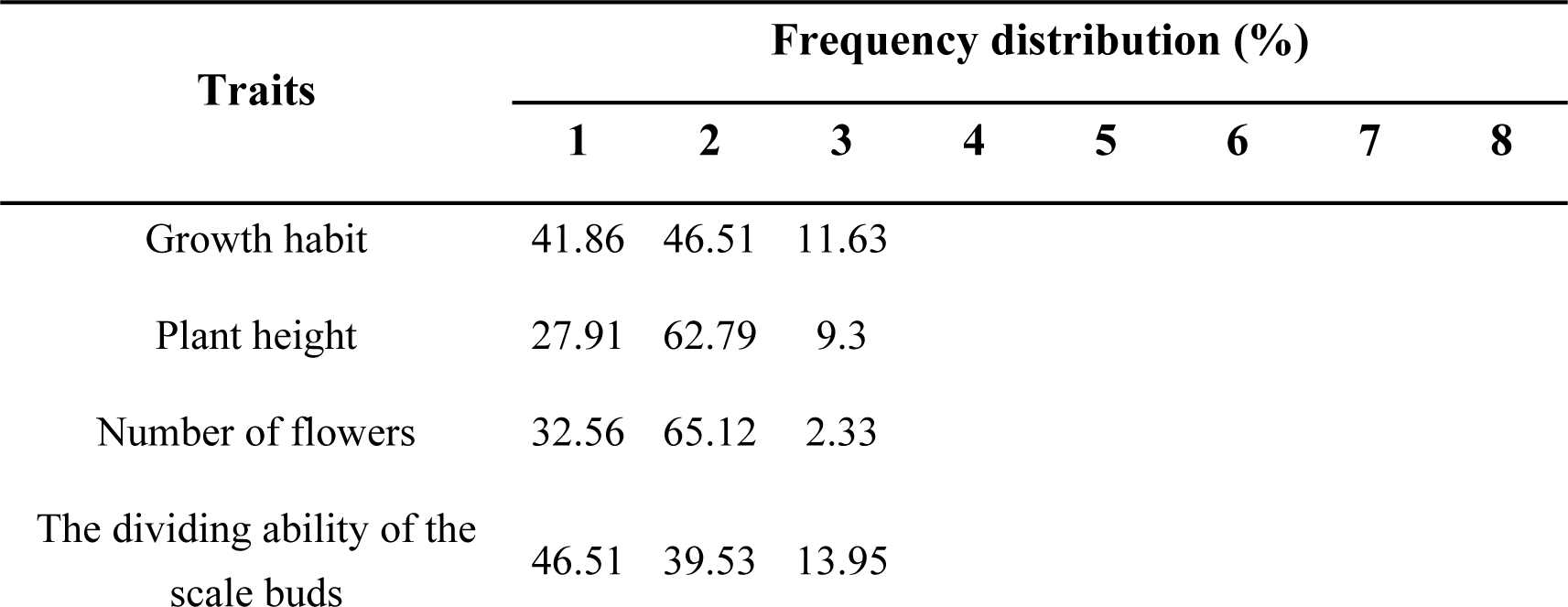
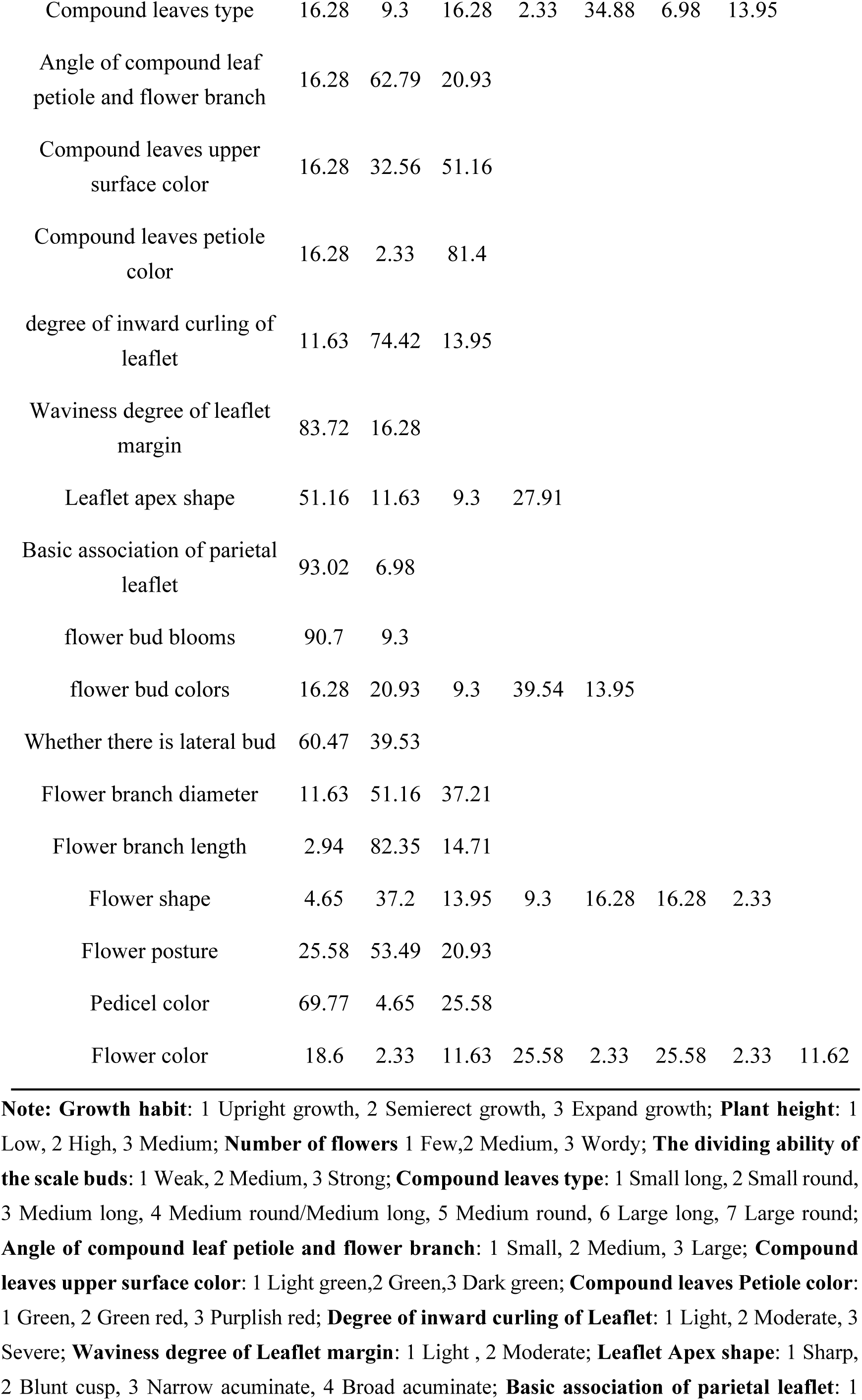
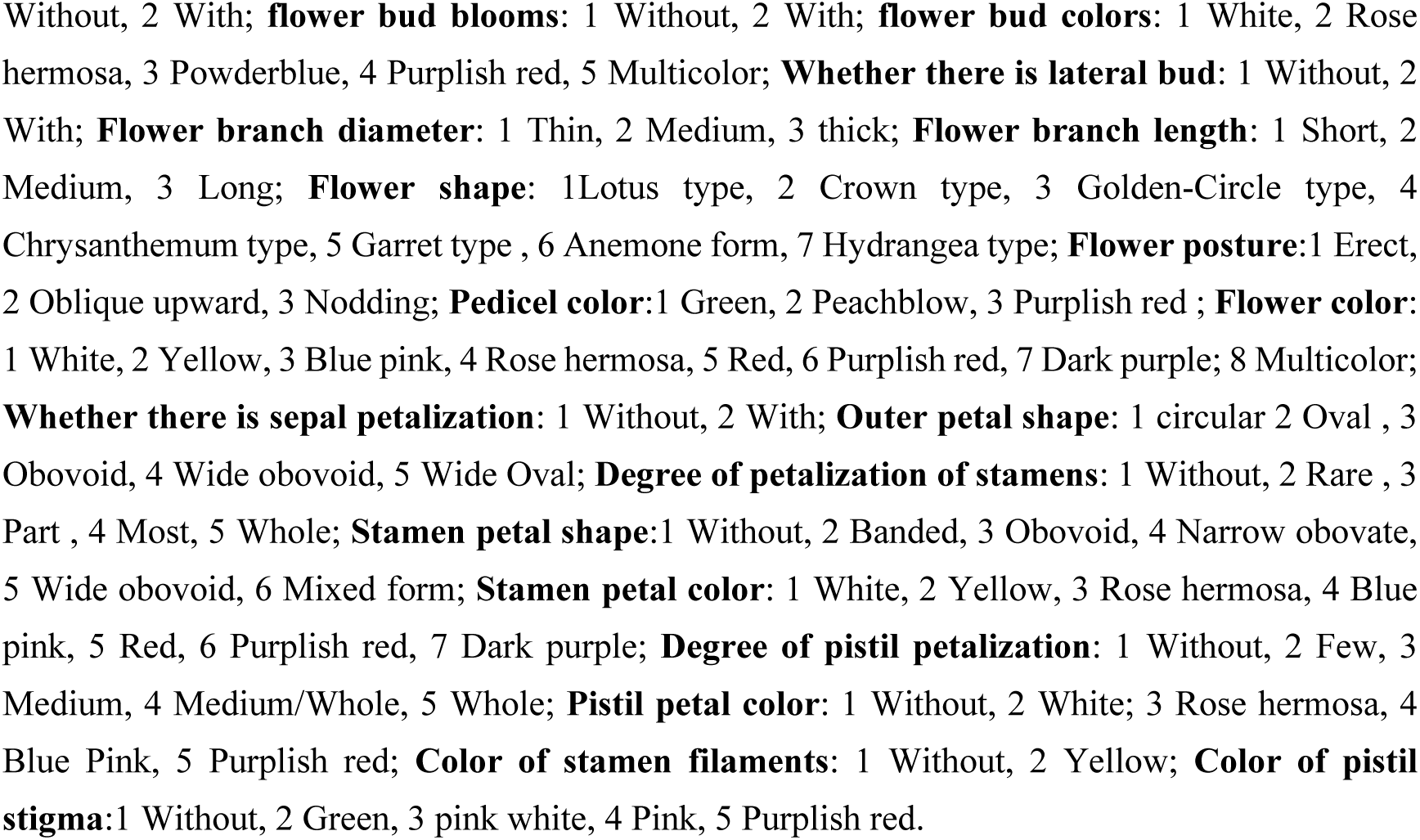
Frequency distribution of peonies of Sect. *Paeonia* qualitative traits.

The coefficient of variation for qualitative traits of 43 plant variety samples ranges from 24.096% to 84.014% (average = 41.933%). The variation degree from large to small was as follows: degree of pistil petalization>color of pistil stigma>pistil petal color>stamen petal color>leaflet apex shape>pedicel color> flower color> compound leaves type> flower shape> outer petal shape> flower bud colors> the dividing ability of the scale buds> growth habit> degree of petalization of stamens> whether there is lateral bud> flower posture> stamen petal shape> plant height> waviness degree of leaflet margin> compound leaves upper surface color> color of stamen filaments> flower branch length> number of flowers> angle of compound leaf petiole and flower branch> flower branch diameter> compound leaves petiole color> flower bud blooms> whether there is sepal petalization> degree of inward curling of leaflet> basic association of parietal leaflet. Among these qualitative traits, 7 indicators had a coefficient of variation exceeding 50%, this indicated that the apparent characteristics of peony samples were relatively complex, and the coefficient of variation of the Degree of pistil petalization was the highest (84.014%), the coefficient of basic association of parietal leaflet was the lowest (24.096).

Descriptive statistical analysis was conducted on 7 quantitative traits of 43 varieties plants of Sect. *Paeonia* samples, which was shown in Table 4. *H’* of quantitative traits ranged from 0.103 to 0.395, with an average of 0.222. The coefficient of variation of the quantitative traits ranged from 11.174% to 42.109%, with an average of 25.753%. Compared to quantitative traits related to flowers, the *H’* of quantitative traits related to whole plants was higher. The *H’* of main stem height was the highest (0.395), which the coefficient of variation was 22.335%, and the number of flower colors was the lowest (0.103), but the coefficient of variation of this trait was relatively high (33.641%). The *H* ’ from high to low was as follows: main stem height> main branch length> flower diameter > number of branches> main branch diameter> crown diameter> number of flower colors. The coefficient of variation from high to low was as follows: number of branches (42.109%)> number of flower colors> crown diameter> main branch length> main stem height> main branch diameter> flower diameter.

**Table 4.**
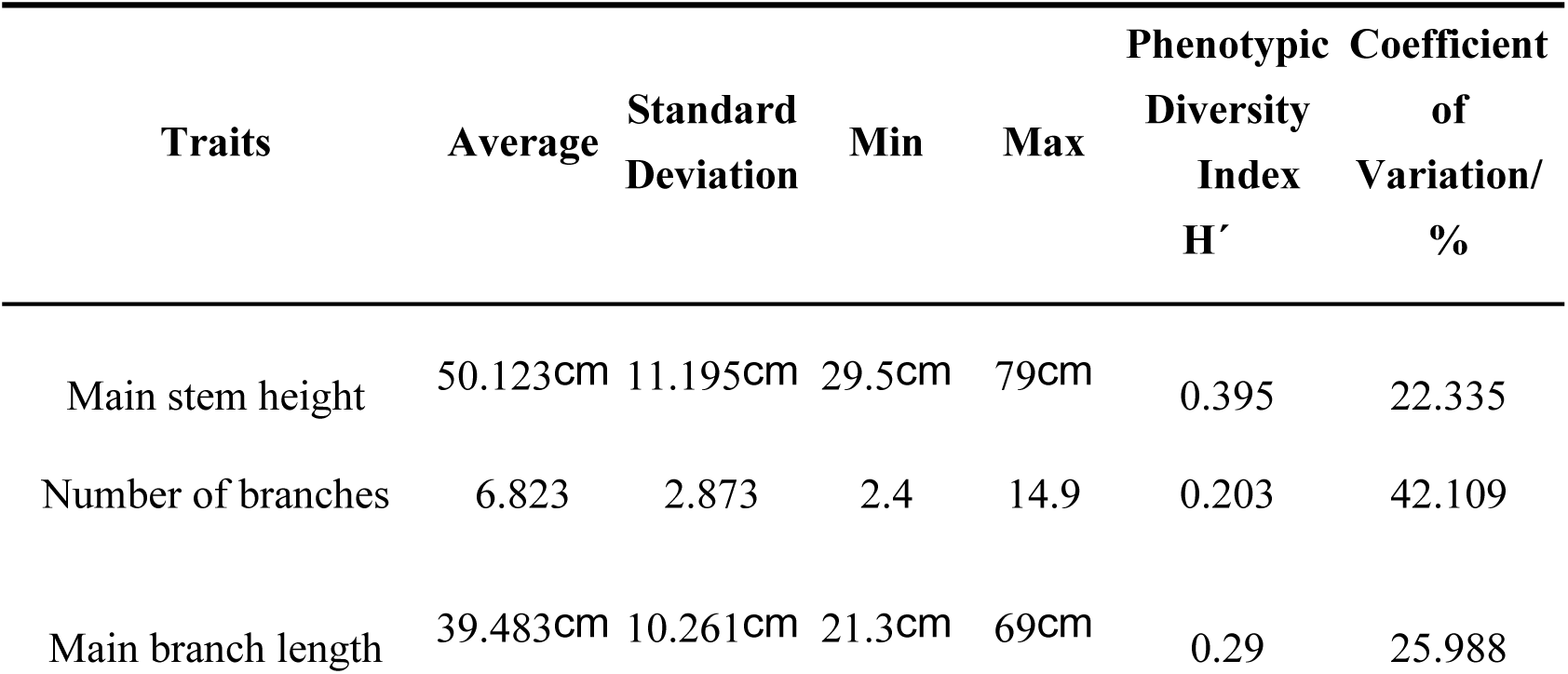
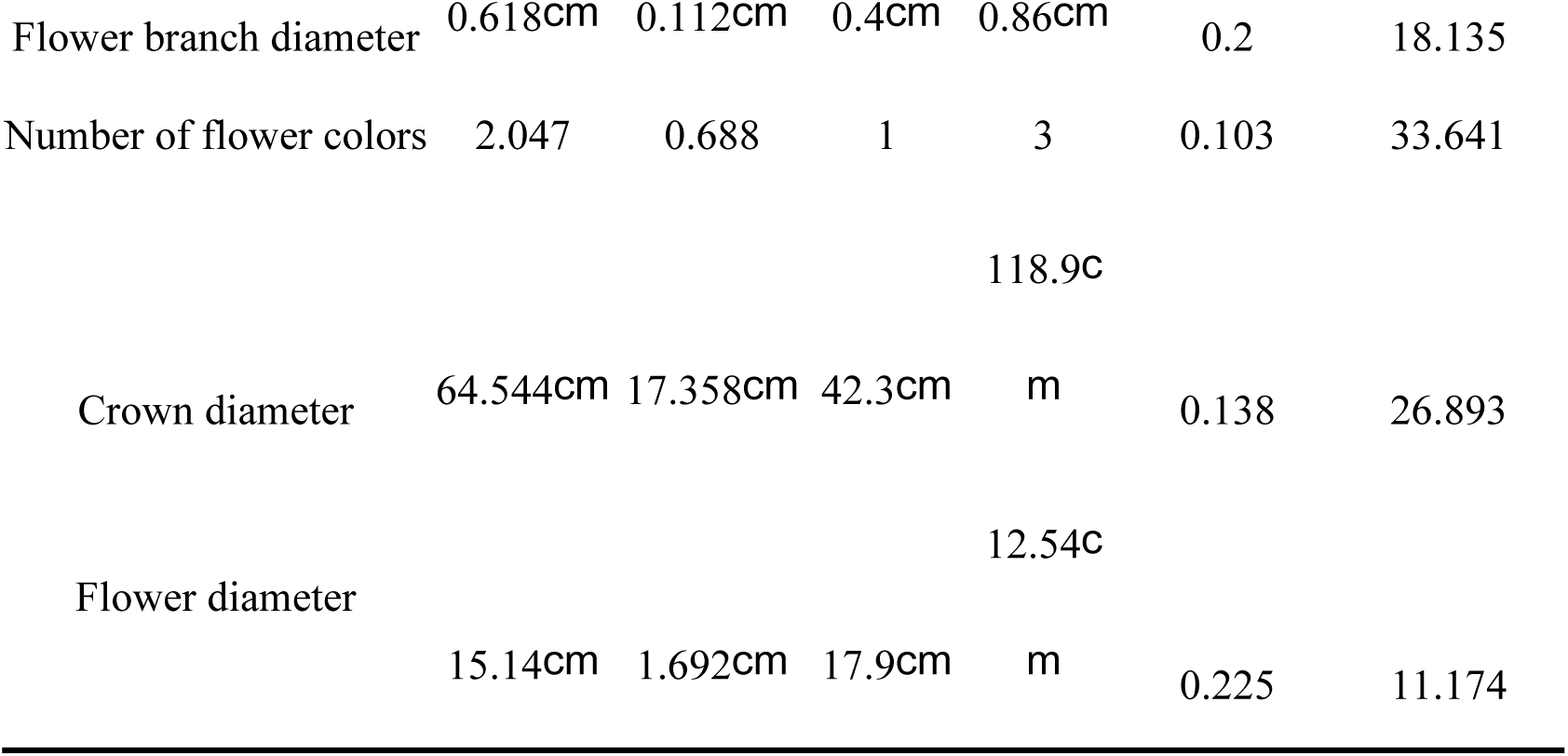
Analysis of diversity of quantitative characters of peonies of Sect. *Paeonia*.

### Principal component analysis of phenotypic traits

Principal component analysis of phenotypic traits is a comprehensive indicator that can fully reflect the role of various phenotypic traits in the composition of diversity. Principal component analysis was conducted on 37 phenotypic traits of tested peonies of Sect. *Paeonia*. We performed the Kaiser–Meyer–Olkin (KMO) test and Bartlett’s spherical test, the KMO was 0.328 and Sig was <0.05, which suggested that the quantitative traits supported PCA. Based on the principle that the characteristic value was greater than 1.0, the eigenvalues of 12 principal components exceeded 1.0 (Table 5), and the cumulative contribution rate of the first 12 extracted principal components reached 78.764%, they reflected the most of information on all indicators and reflected the proportion of various phenotypic traits in the principal components of tested varieties.

**Table 5.** Principal component analysis results of 37 phenotypic traits in 43 varieties of peonies of Sect. *Paeonia* samples

| Traits | Principal component |  |  |  |  |  |  |  |  |  |  |  |
| --- | --- | --- | --- | --- | --- | --- | --- | --- | --- | --- | --- | --- |
|  | 1 | 2 | 3 | 4 | 5 | 6 | 7 | 8 | 9 | 10 | 11 | 12 |
| Growth habit | 0.271 | -0.452 | 0.016 | -0.232 | 0.282 | 0.027 | -0.396 | 0.123 | -0.335 | 0.201 | 0.157 | -0.046 |
| Plant height | 0.884 | 0.081 | -0.1 | -0.084 | -0.02 | 0.011 | 0.089 | 0.095 | -0.009 | 0.192 | -0.173 | 0.01 |
| Number of flowers | 0.65 | 0.14 | 0.069 | 0.061 | -0.239 | 0.293 | -0.2 | -0.014 | 0.082 | 0.117 | 0.204 | 0.204 |
| The dividing ability of the scale buds | 0.498 | 0.556 | -0.355 | 0.083 | 0.042 | 0.246 | -0.049 | -0.143 | 0.118 | 0.231 | 0.015 | 0.175 |
| Compound leaves type | 0.115 | -0.308 | -0.293 | -0.006 | 0.167 | 0.029 | -0.093 | 0.705 | 0.056 | 0.237 | 0.02 | 0.062 |
| Angle of compound leaf petiole and flower branch | 0.198 | -0.217 | -0.14 | 0.414 | 0.16 | 0.148 | 0.517 | -0.082 | -0.272 | -0.156 | 0.298 | -0.014 |
| Compound leaves upper surface color | -0.541 | -0.166 | 0.005 | -0.168 | 0.101 | -0.055 | 0.357 | 0.228 | 0.15 | 0.357 | 0.222 | -0.093 |
| Degree of inward curling of leaflet | -0.029 | 0.428 | 0.149 | 0.275 | 0.439 | -0.282 | 0.231 | -0.18 | 0.071 | 0.094 | -0.169 | 0.152 |
| Waviness degree of leaflet margin | 0.164 | 0.237 | 0.376 | 0.446 | -0.015 | 0.433 | 0.252 | -0.096 | 0.041 | 0.309 | -0.162 | 0.001 |
| Leaflet apex shape | 0.235 | -0.152 | -0.041 | -0.316 | 0.134 | -0.125 | 0.534 | 0.068 | 0.196 | 0.023 | -0.27 | 0.215 |
| Basic association of parietal leaflet | 0.19 | 0.377 | -0.383 | 0.194 | -0.049 | -0.01 | 0.005 | -0.348 | 0.362 | 0.284 | 0.156 | -0.096 |
| Flower bud blooms | 0.064 | 0.239 | -0.348 | 0.472 | -0.248 | 0.021 | 0.171 | 0.4 | -0.088 | -0.011 | 0.219 | 0.311 |
| Flower bud colors | 0.144 | 0.671 | 0.312 | 0.09 | -0.005 | 0.147 | 0.02 | 0.031 | 0.026 | -0.145 | 0.15 | -0.022 |
| Whether there is lateral bud | 0.19 | -0.113 | 0.129 | 0.134 | -0.068 | 0.316 | 0.156 | -0.007 | -0.686 | 0.179 | -0.303 | 0.09 |
| Flower branch diameter | 0.235 | -0.444 | -0.005 | 0.142 | -0.158 | 0.214 | -0.168 | 0.224 | 0.027 | -0.418 | -0.232 | 0.013 |
| Flower branch length | 0.883 | 0.154 | -0.054 | 0.102 | 0.032 | -0.081 | 0.065 | 0.18 | -0.113 | 0.158 | -0.026 | -0.032 |
| Flower posture | 0.448 | -0.503 | -0.329 | 0.104 | 0.401 | -0.045 | -0.188 | -0.132 | -0.053 | 0.073 | 0.109 | -0.083 |
| Flower diameter | 0.286 | -0.647 | 0.072 | 0.037 | 0.119 | -0.175 | 0.17 | 0.009 | 0.125 | -0.169 | 0.423 | 0.077 |
| Flower shape | 0.029 | -0.049 | 0.568 | -0.398 | 0.067 | 0.156 | 0.052 | 0.124 | 0.124 | 0.088 | -0.076 | 0.516 |
| Number of flower colors | 0.081 | 0.203 | 0.542 | 0.027 | 0.045 | -0.476 | -0.232 | 0.01 | -0.086 | -0.187 | 0.122 | 0.386 |
| Flower color | -0.009 | 0.562 | 0.528 | -0.269 | 0.052 | 0.299 | -0.115 | 0.084 | 0.047 | 0.085 | 0.256 | -0.126 |
| Whether there is sepal petalization | 0.343 | 0.209 | 0.075 | 0.101 | 0.516 | 0.271 | 0.126 | 0.227 | 0.011 | -0.289 | 0.09 | -0.117 |
| Outer petal shape | -0.439 | -0.033 | -0.233 | -0.008 | 0.409 | 0.076 | -0.242 | 0.206 | 0.235 | 0.31 | -0.068 | 0.17 |
| Degree of petalization of stamens | 0.12 | -0.684 | 0.351 | -0.179 | -0.251 | 0.199 | 0.12 | -0.137 | 0.039 | 0.277 | 0.178 | -0.026 |
| Stamen petal shape | -0.05 | 0.009 | -0.016 | -0.345 | 0.449 | 0.098 | 0.358 | -0.083 | 0.192 | -0.206 | 0.038 | 0.023 |
| Stamen petal color | -0.1 | 0.611 | 0.146 | -0.227 | 0.064 | 0.238 | -0.043 | 0.321 | 0.083 | 0.035 | -0.006 | -0.359 |
| Degree of pistil petalization | 0.201 | -0.532 | 0.221 | 0.239 | 0.509 | 0.07 | -0.094 | -0.082 | 0.047 | 0.054 | -0.212 | -0.028 |
| Pistil petal color | 0.162 | -0.168 | 0.57 | 0.514 | 0.368 | 0.059 | -0.213 | -0.056 | 0.089 | 0.03 | 0.008 | -0.164 |
| Compound leaves petiole color | 0.069 | 0.284 | 0.079 | -0.117 | 0.08 | -0.654 | 0.014 | -0.076 | -0.315 | 0.396 | -0.041 | -0.08 |
| Pedicel color | 0.261 | 0.357 | 0.005 | -0.502 | 0.093 | -0.053 | 0.283 | 0.106 | -0.363 | -0.097 | 0.031 | -0.222 |
| Color of stamen filaments | -0.064 | 0.731 | -0.233 | 0.168 | 0.272 | -0.2 | -0.164 | 0.147 | -0.026 | -0.223 | -0.168 | 0.034 |
| Color of pistil stigma | 0.233 | 0.084 | 0.383 | 0.437 | -0.262 | -0.429 | 0.143 | 0.367 | 0.105 | 0.057 | 0.091 | -0.119 |
| Main stem height | 0.881 | -0.046 | 0.136 | -0.126 | -0.053 | -0.261 | -0.027 | -0.029 | 0.172 | -0.033 | -0.018 | -0.076 |
| Crown diameter | 0.853 | 0.042 | -0.148 | -0.139 | 0.071 | -0.079 | 0.039 | -0.084 | -0.021 | -0.113 | 0.124 | 0.061 |
| Main branch length | 0.888 | -0.001 | 0.111 | -0.097 | -0.056 | -0.175 | -0.069 | 0.003 | 0.113 | 0.049 | -0.002 | -0.145 |
| Main branch diameter | 0.547 | -0.321 | 0.025 | -0.026 | -0.282 | -0.022 | 0.09 | 0.112 | 0.35 | -0.097 | -0.368 | -0.189 |
| Number of branches | 0.619 | 0.281 | -0.212 | -0.317 | 0.051 | 0.221 | -0.16 | -0.141 | -0.053 | -0.105 | 0.072 | 0.187 |
| Eigenvalue | 6.686 | 5.019 | 2.691 | 2.372 | 2.041 | 2.006 | 1.674 | 1.497 | 1.466 | 1.435 | 1.167 | 1.088 |
| Contribution /% | 18.071 | 13.565 | 7.272 | 6.411 | 5.517 | 5.421 | 4.525 | 4.046 | 3.962 | 3.879 | 3.155 | 2.941 |
| Cumulative contribution /% | 18.071 | 31.636 | 38.908 | 45.319 | 50.836 | 56.257 | 60.781 | 64.827 | 68.79 | 72.669 | 75.823 | 78.764 |

The contribution of the first principal component was 18.071%, and the eigenvalue was 6.686. The top 5 characteristic vectors with the highest absolute value were all positively correlatedwere, plant height, flower branch length, main stem height, crown diameter and main branch length were the main phenotypic traits that affect the principal components, their featurevectors were respectively: 0.884, 0.883, 0.881, 0.853, 0.888. The first principal component was mainly related to the overall height of the plant and branch.

The contribution of the second principal component was 13.565%, and the eigenvalue was 5.019. The traits with higher absolute values of featurevector include color of stamen filaments, flower bud colors, stamen petal color, flower diameter, and degree of petalization of stamens, and their featurevectors were respectively: 0.731, 0.671, 0.611, -0.647, -0.684. The featurevectors of principal component 2 were mostly related to stamens.

The contribution of the third principal component was 7.272%, and the eigenvalue was 2.691. Pistil petal color, flower shape, number of flower colors, flower color, color of pistil stigma and basic association of parietal leaflet were the main indexs, and their featurevectors were respectively: 0.570, 0.568, 0.542, 0.528, 0.383, -0.383. The featurevectors of principal component 3 were mainly related to the color of flowers and pistils.

The contribution of the fourth principal component was 6.411%, and the eigenvalue was 2.372. The traits with higher absolute values of featurevectors were pistil petal color, flower bud blooms, waviness degree of leaflet margin, color of pistil stigma, pedicel color, and their featurevectors were respectively: 0.514, 0.472, 0.446, 0.437, - 0.502. The featurevectors of principal component 4 were mostly related to the characteristics of the flowers.

The contribution of the fifth principal component was 5.517%, and the eigenvalue was 2.041. The traits with higher absolute values of featurevector include whether there was sepal petalization, degree of pistil petalizatio, stamen petal shape, degree of inward curling of leaflet, outer petal shape, and their featurevectors were respectively: 0.516, 0.509, 0.449, 0.439, 0.409. The above traits were positively correlated with the principal component 5, which mainly related to the petalization of sepals and stamens.

The contribution of the sixth principal component was 5.421%, and the eigenvalue was 2.006. The top 5 characteristic vectors with the highest absolute value were: waviness degree of leaflet margin, whether there is lateral bud color of pistil stigma, number of flower colors and compound leaves petiole color, and their featurevectors were respectively: 0.433, 0.316, 0.429, -0.476, -0.654. Principal component 5 had a high correlation with color traits.

The contribution of the seventh principal component was 4.525%, and the eigenvalue was 1.674. The traits with higher absolute values of featurevectors were: leaflet apex shape, angle of compound leaf petiole and flower branch, stamen petal shape, compound leaves upper surface color and growth habit, and their featurevectors were respectively: 0.534, 0.517, 0.357, 0.358, -0.396. This principal component was mainly related to leaf traits.

The contribution of the eighth principal component was 4.046%, and the eigenvalue was 1.497. The traits with higher absolute values of featurevector include compound leaves type, flower bud blooms, color of pistil stigma, stamen petal color and basic association of parietal leaflet, and their featurevectors were respectively: 0.705, 0.4000, 0.376, 0.321, -0.348.

The contribution of the ninth principal component was 3.962%, and the eigenvalue was 1.466. The traits with higher absolute values of featurevectors were: basic association of parietal leaflet, main branch diameter, growth habit, pedicel color, whether there is lateral bud, and their featurevectors were respectively: 0.362, 0.350, - 0.335, -0.363, -0.686.

The contribution of the tenth principal component was 3.879%, and the eigenvalue was 1.435. The traits with higher absolute values of featurevector include compound leaves petiole color, compound leaves upper surface color, outer petal shape, waviness degree of leaflet margin, flower branch diameter, and their featurevectors were respectively: 0.396, 0.357, 0.310, 0.309, -0.418. The main positively correlated traits were related to leaf color and shape.

The contribution of the eleventh principal component was 3.155%, and the eigenvalue was 1.167. The traits with higher absolute values of featurevector include: flower diameter, angle of compound leaf petiole and flower branch, leaflet apex shape, whether there is lateral bud, main branch diameter, and their featurevectors were respectively: 0.423, 0.298, -0.270, -0.303, -0.368. The eleventh principal component mainly reflected the quantitative traits related to flower and whole plant.

The contribution of the twelfth principal component was 2.941%, and the eigenvalue was 1.088. The traits with higher absolute values of featurevector include: flower shape, number of flower colors, flower bud blooms, pedicel color, stamen petal color, and their featurevectors were respectively: 0.516, 0.386, 0.311, -0.222, -0.359. The principal component 12 mainly reflected traits related to flower.

Based on the comprehensive analysis of contribution and eigenvalues, waviness degree of leaflet margin, basic association of parietal leaflet, flower bud blooms, whether there is lateral bud, number of flower colors, stamen petal color, pedicel color, color of pistil stigma, were selected from the principal components as important indicators for evaluating the phenotypic genetic diversity of plants of Sect. *Paeonia* samples.

### Comprehensive evaluation of phenotypic traits

Comprehensive evaluation of 43 germplasm samples used the score coefficients of each principal component obtained from principal component analysis. The linear equations of F_1_, F_2_… F_12_ were obtained based on 12 principal component coefficients (TableS1). Substituted the standardized data of 37 phenotypic traits into the 12 linear equations mentioned above to obtain the scores of principal components for each peony of Sect. *Paeonia* variety (Table 6). F_i_ represented the score of the i-th principal component, and X_1_, X_2_, X_3_ . . . . . . X_37_ represent the standardized data of each phenotypic trait.

**Table 6.**
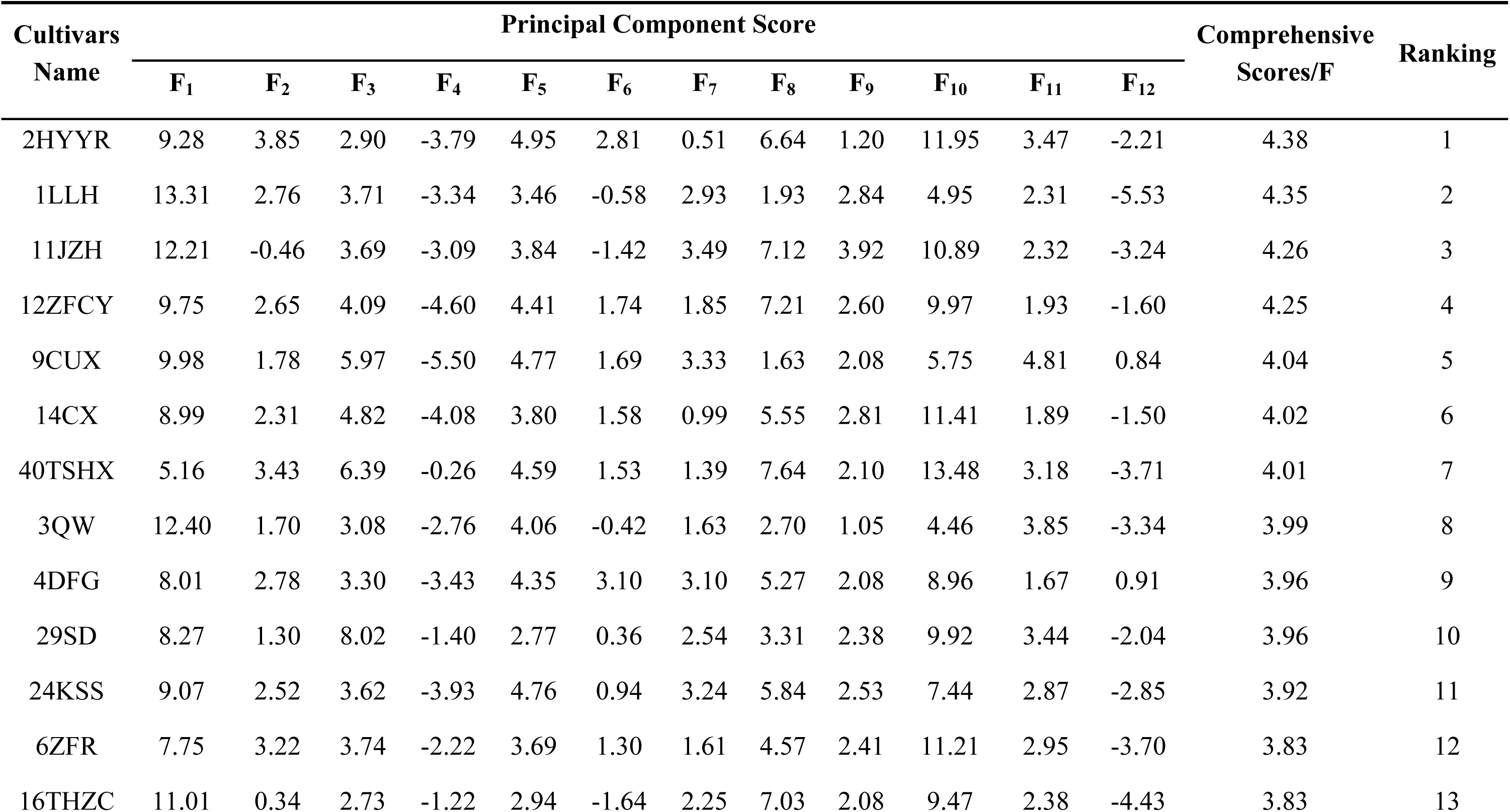
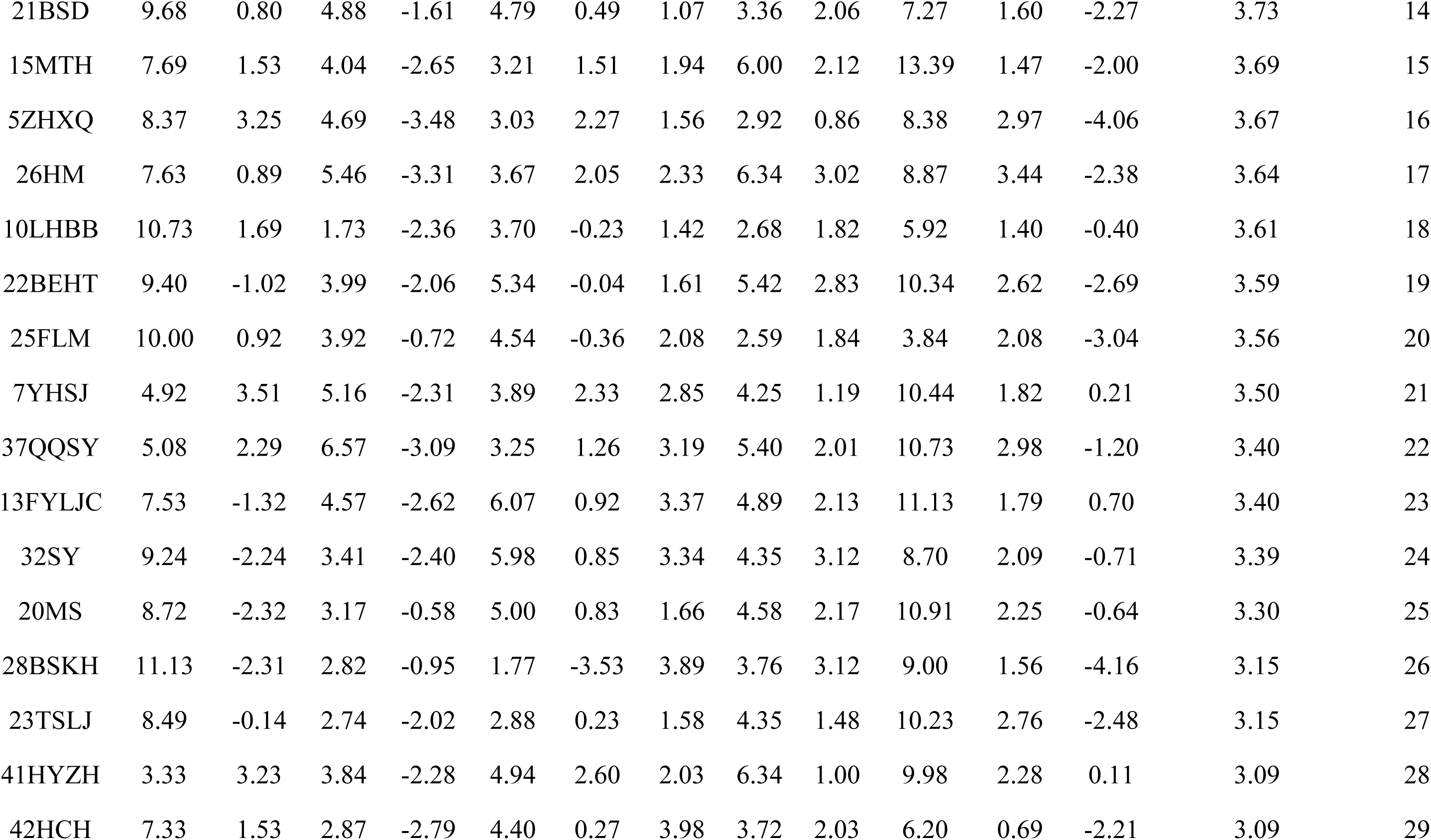
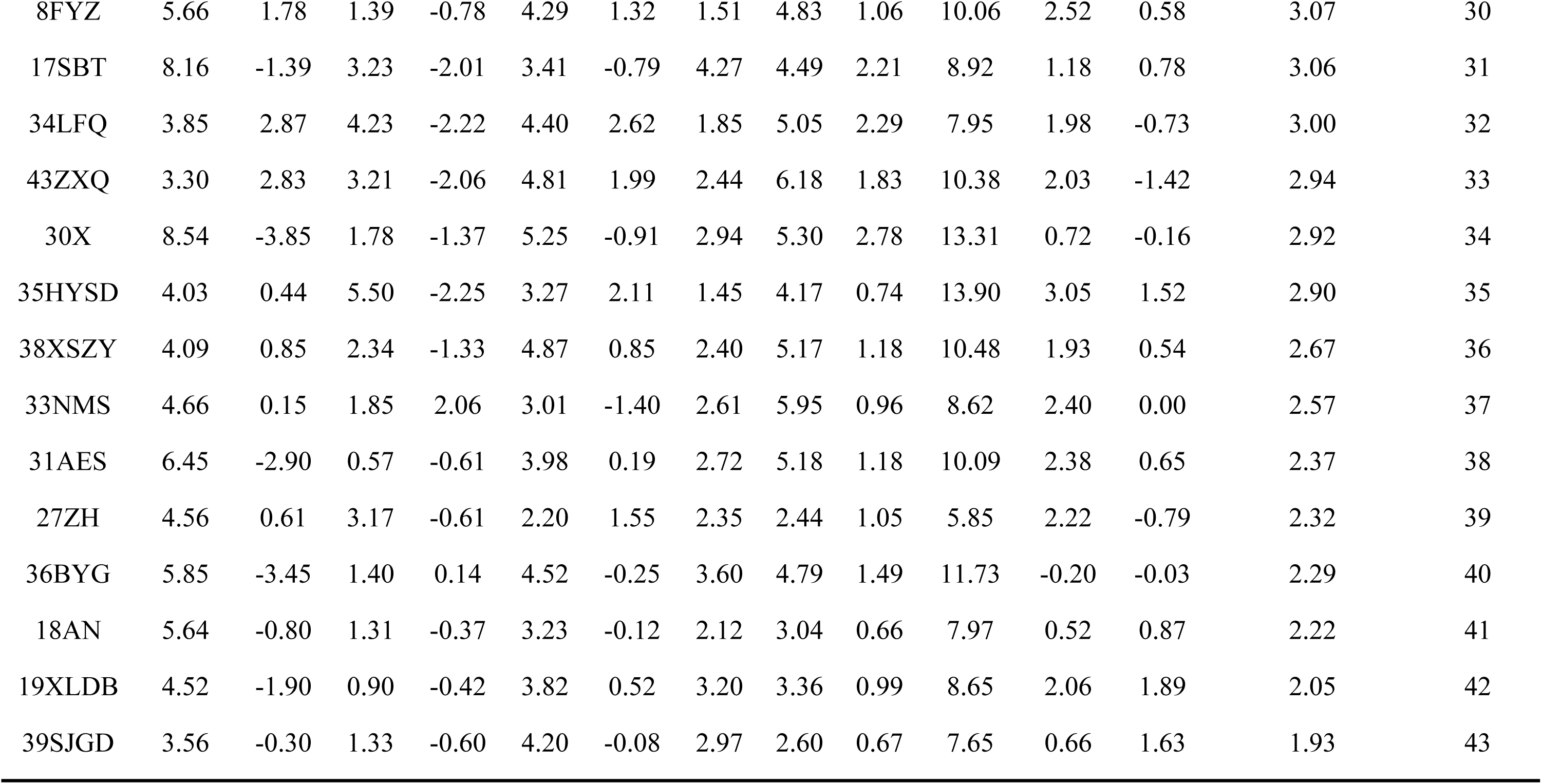
Comprehensive evaluation about phenotypic traits of peonies of Sect. *Paeonia*

In addition, a comprehensive evaluation model for phenotypic diversity was constructed based on 12 principal component coefficients and the contribution of each principal component:

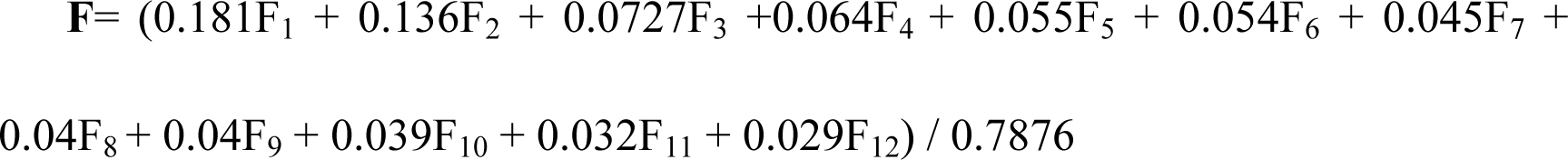

Calculated the comprehensive score of each peony variety of Sect. *Paeonia* based on the contribution of 12 principal components, and comprehensively evaluated their phenotypic diversity. The comprehensive scores/F value of the 43 varieties ranged from 1.93 to 4.38. The top 5 varieties with the highest F value were 2HYYR (4.38), 1LLH (4.35), 11JZH (4.26), 12ZFCY (4.25), 9CUX (4.40). The above 5 varieties have relatively high comprehensive scores, and their phenotypic traits can effectively represent the diversity of phenotypic traits among multiple varieties. This indicated that during the cultivation and hybridization process, these varieties of peony plants have inherited richer traits.

### Cluster analysis

Cluster analysis was conducted on the germplasm resources and phenotypic traits of peonie of Sect. *Paeonia*, and the results of cluster analysis are shown in Figure 2. Figure 2(A) showed the clustering analysis of 37 phenotypic traits, while Figure 2(B) showed the clustering analysis of 43 germplasm resources.

**Figure 2.**
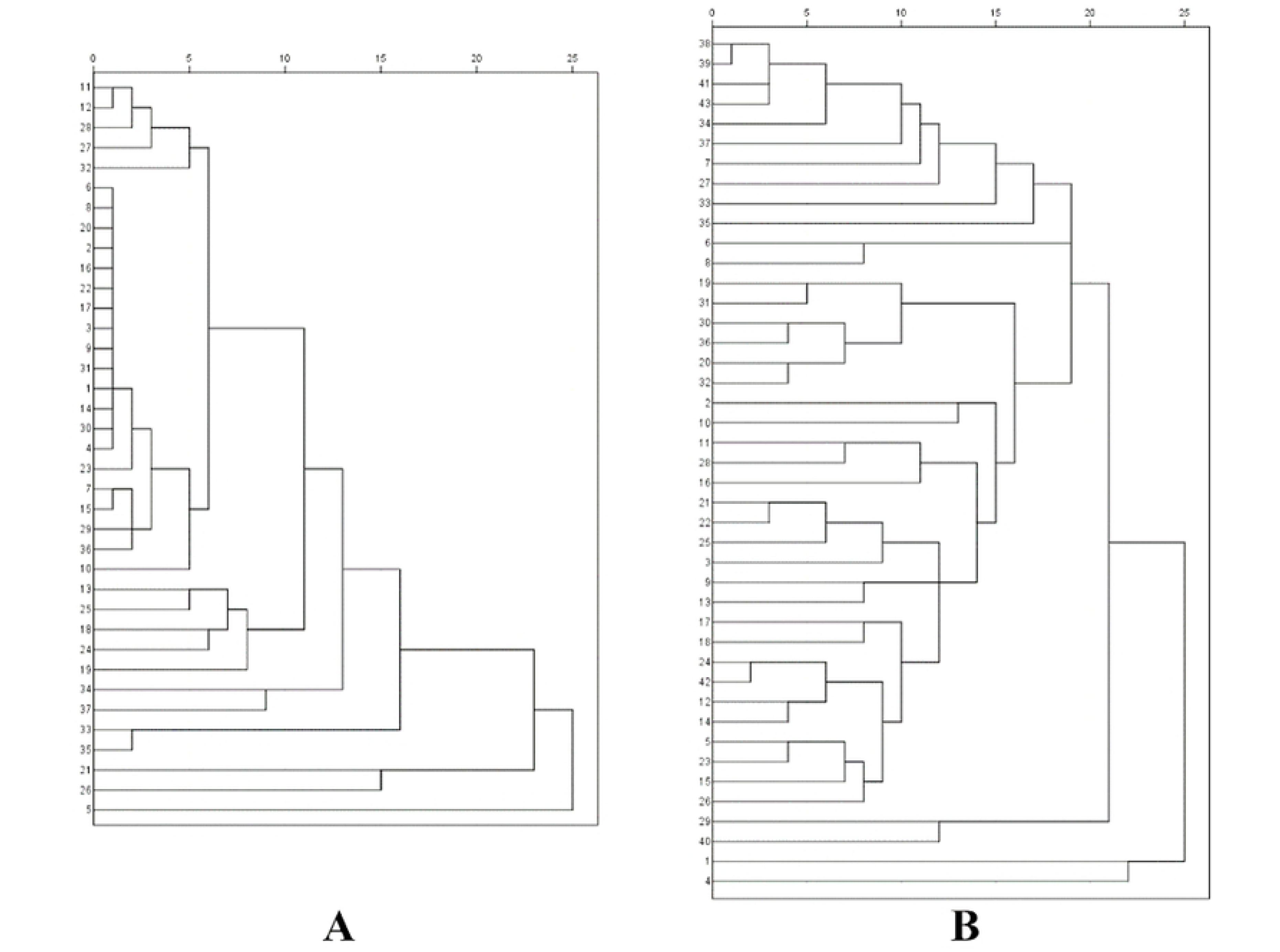
(A) Clustering figure based on phenotypic charactors. (B) Clustering figure based on cultivars

The squared *Euclidean distance* was considered the genetic distance. All phenotypic traits were clustered into 4 groups at a genetic distance of 15 (Figure 2). The first group contained 32 phenotypic traits. The second group contained 2 phenotypic traits, which was main stem height and main branch length, which indicated there was a direct correlation between the height of plants and the height of the main stem. The third group contains 2 phenotypic traits, which was flower color and stamen petal color, the correlation between these two colors related traits was stronger. The fourth group contains 1 phenotypic trait, which was compound leaves type, Indicated a low degree of association between the type of compound leaves and other phenotypic traits.The overall results indicated that most of the indicators selected in this study are highly correlated, and the evaluation of these indicators could effectively determine the genetic diversity of peony plants. This further indicated that the evaluation indicators in the Agricultural Industry Standard of the People’s Republic of China (NY/T2225-2012-Guidelines for Specificity, Consistency and Stability Testing of New Plant Varieties were scientific and effective.

All varieties of peony germplasm resources were clustered into 4 groups at a *Euclidean distance* of 20 (Figure 2). The second group included two varieties, which were 29SD and 40TSHX. The third group only had one variety, which was 1LLH, while the fourth group only had one variety, which was 4DFG. The other 39 varieties were all clustered in the first group. This indicated that the richness of phenotypic traits in the tested samples we selected were very close. These hybrid varieties of peony plantsnot only had different flower colors, but also exhibited various morphological changes in petal layering, leaves, and plant morphology. The genetic diversity was rich, which was an optimistic result for ornamental plants.

## Discussion

Plant phenotypic diversity is a comprehensive reflection of genetics and environment [30,31]. Mathematical statistics are commonly used to visually display the degree of genetic diversity and variation in plant populations, which is also a common method for studying plant epigenetic morphology. This study analyzed the genetic diversity of 37 phenotypic traits from 43 ornamental peony varieties, and comprehensively analyzed the results of *H’* [32] and coefficient of variation [33] to understand the variation of germplasm resources from multiple aspects and explore the breeding value of each ornamental peony variety. The comprehensive evaluation of genetic diversity can evaluate the phenotypic richness of species populations from a holistic perspective. Among the traits involved in this study, the highest H’ was compound leaves type, which are a key structure for the stress resistance of peony. Although compound leaves do not have a large total leaf area, their resistance to wind and rain is relatively reduced, making them an evolutionary feature that adapts to the environment. Therefore, the regulation and improvement of compound leaf types is a way to improve environmental adaptability [34]. Due to differences in plant morphology, height and habits, peony also exhibits a rich variety of compound leaves types, which are key features for adapting to the environment and enhancing its stress resistance. In addition, the color of the flowers and the degree of petalization of the stamens have a relatively high H ’value.

The complex color of flowers is the main characteristic of peony plants of Sect *Paeonia.*, which gives them great ornamental value. The color grading of peony flowers is very complex, and some varieties exhibit compound colors. The degree of petalization of stamens directly affects the morphology of flowers. The key indicator for the natural transformation of a single petal flower into a double petal flower is stamen petalization. Plant varieties with different petal layers can be cultivated by artificially inducing the degree of stamen petalization. When the degree of petalization of stamens is high and the degree of petalization of pistils is low, it will show a high and clustered flower shape, for example, the crown shaped flower is a typical variety. Therefore, traits with higher H ’values will affect the macroscopic morphology of peony, resulting in significant differences among different varieties. The H’ of the degree of pistil petalization in this study is only 0.043, and 76.74% of the varieties here showed without pistil petalization. This indicated that in the process of artificial cultivation of peony varieties, the complex flower pattern is mainly transformed into a double petal shape by abandoning the stamens, but the situation of abandoning the pistils is relatively rare.

Principal component analysis [35,36] is a comprehensive evaluation of data by calculating the weight coefficients of different variables. By integrating a large number of variables into fewer variables, the data is easier to analyze intuitively.Principal component analysis is particularly suitable for screening and evaluating excellent plant varieties. This study integrated 37 phenotypic traits into 12 principal components through principal component analysis. By comparing the contribution rates of principal components, we found that the contribution rates of principal components 1 and 2 were much higher than those of the other 10 principal components, which were 18.071% and 13.565%, respectively. The main traits that affected these two principal components were quantitative traits, color traits, and degree of stamen petalization. This also indicates that the phenotypic traits of different varieties of peony plants undergo the most diverse changes during the continuous domestication and hybridization process, including the appearance height of the plant, the color of flowers and other organs, and the changes in flower shape resulting from the degree of petalization of stamens. This result also corresponds to the result of H’, where the degree of petalization of stamens has a significant impact on the morphology of peony plants, providing a favorable reference for the breeding of new varieties. In fact, the changes in plant height, growth habits, and leaf shape are important manifestations of adaptation to the environment [37]. For example, for upright plants with higher plant height, the types of compound leaves are mostly small long or medium round. This is to reduce the stress on tall plants in extreme weather conditions such as wind and rain, and enhance their stress resistance. The highest value of H ’in quantitative traits was main stem height, while the lowest value was number of flower colors. The phenotypic traits are relatively more affected by annual environment and climate change, and different varieties have different adaptability to climate change. The phenotypic traits of the same variety may also vary in different years, depending on environmental factors and plant age. This study only recorded data and information for two years, and since the growth environment of all plant samples in this study was consistent, it may not be possible to comprehensively display all the epigenetic features. Especially quantitative traits are easily influenced by environmental changes, resulting in significant differences in variation. However, comparing different varieties under the same living environment has significant advantages, as we can explore the characteristics of the superior varieties through this study, and identify key processes that can be regulated and intervened in the cultivation process of new varieties through extensive data analysis, such as degree of petalization of stamens. Flower color is the main characteristic that determines the ornamental value of peony, and multicolored flower have more ornamental value. Therefore, in recent years, many researchers have focused on exploring the relevant genes and enzymes that regulate peony flower color. For example, pelargonidin 3-O-glucoside, cyanidin-3,5-di-O-glucoside, and peonidin-3,5-di-O-glucoside are important compounds that regulate the color of peony flowers [38].

The comprehensive evaluation scheme can effectively compare the differences between different plant species [39]. Comprehensive evaluation is a comprehensive score obtained by normalizing the relevant parameters of principal component analysis [40]. The comprehensive score of all peony varieties in this study was positive, this indicates that the population of plants of Sect *Paeonia* is relatively abundant in terms of genetic diversity performance. There were 7 varieties with a comprehensive score higher than 4, all of which were Central Plains varieties. In addition, this study conducted cluster analysis using variety and phenotypic traits as variables, respectively. The color of the stamen petal and the color of the flower cluster together, indicating that the degree of stamen petal will to some extent affect the overall color realization of the flower. In terms of colors traits, we can observe that the color of stamen petal petals sometimes does not match the color of the flower. Some of these become part of the multicolored flower, while others are due to the low degree of stamen petal petals when the flower is fully open, making its color not obvious. In the clustering of varieties, we did not find any obvious distinguishing features. For example, Dutch varieties and Chinese varieties were not clearly clustered separately, indicating that the source of germplasm resources is not the key to determining the performance of their apparent traits, while environmental regulation and artificial intervention seem to be more important.

## Conclusion

The 43 varieties of plant samples of Sect *Paeonia* collected in this study showed rich phenotypic diversity changes after research. We found that the traits that significantly regulate the appearance differences of different varieties of peony include the degree of stamen petalization, plant height, and compound leaves type. Of course, other phenotypic traits have also contributed to enriching the phenotypic diversity of peony resources. The flowering period of peony is very short, so the number of varieties collected in this experiment cannot represent all the planting in the local area. Therefore, in future research, the investigation of germplasm resources will be expanded. At the same time, it is necessary to expand the scope of the investigation environment and explore the genetic diversity performance of the same variety in different planting environments, which will be conducive to exploring the impact of environmental factors on genetic performance. Overall, this study provided a reference for the genetic breeding of ornamental peony. In order to cultivate new varieties of peony with more obvious phenotypic diversity, the effect of changing plant appearance can be achieved by regulating phenotypic traits such as flower color, degree of stamen petalization, and color of stamen petalization. It is also possible to improve stress resistance by adjusting the compound leaves type, plant height, and crown diameter.

## Author contributions

Hui-yan Cao and Shi-yi Xu implemented the research process; Mei-qi Liu, Shan Jiang and Leng-leng Ma helped collect plant materials; Jian-hao Wu, Xiao-Zhuang Zhang, Ling-yang Kong and Wei-chao Ren helped with preliminary data organization; Wei Ma and Xiu-bo Liu guided article writing; Zhi-yang Liu and Xi Chen were assistance during field trips and data analyses.

The data were collected by HYC and SYX. The materials were collected from Germplasm Resource Nursery of Harbin Academy of Agricultural Sciences.

## Supporting information

**S1 Table. The linear equation of 12 principal components.**

## References

1. Xiao PX, Li Y, Lu J, Zuo H, Pingcuo G, Ying H, et al. High-quality assembly and methylome of a Tibetan wild tree peony genome (*Paeonia ludlowii)* reveal the evolution of giant genome architecture. Hortic Res. 2023;10(12):uhad241.

2. Yang L, Wan X, Zhou R, Yuan Y. The Composition and Function of the Rhizosphere Bacterial Community of *Paeonia lactiflora* Varies with the Cultivar. Biology (Basel). 2023;12(11):1363.

3. Cai H, Xu R, Tian P, Zhang M, Zhu L, Yin T, et al. Complete Chloroplast Genomes and the Phylogenetic Analysis of Three Native Species of Paeoniaceae from the Sino-Himalayan Flora Subkingdom. Int J Mol Sci. 2023; 25(1):257.

4. Li P, Shen J, Wang Z, Liu S, Liu Q, Li Y, et al. Genus Paeonia: A comprehensive review on traditional uses, phytochemistry, pharmacological activities, clinical application, and toxicology. J Ethnopharmacol. 2021; 269:113708.

5. Wang Q, Zhu J, Li B, Li S, Yang Y, Wang Q, et al. Functional identification of anthocyanin glucosyltransferase genes: a Ps3GT catalyzes pelargonidin to pelargonidin 3-O-glucoside painting the vivid red flower color of Paeonia. Planta. 2023; 257(4):65.

6. Liu G, Li Y, Sun X, Guo X, Jiang N, Fang Y, et al. Association study of SNP locus for color related traits in herbaceous peony (*Paeonia lactiflora Pall.*) using SLAF-seq. Front Plant Sci. 2022; 13:1032449.

7. Song J, Yang J, Jeong BR. Synergistic Effects of Silicon and Preservative on Promoting Postharvest Performance of Cut Flowers of Peony (*Paeonia lactiflora* Pall.). Int J Mol Sci. 2022; 23(21):13211.

8. Lu YZ, Zhang CQ, Yu BX, Zhang EH, Quan H, Yin X, et al. The seed oil of *Paeonia ludlowii* ameliorates Aβ25-35-induced Alzheimer’s disease in rats. Food Sci Nutr. 2021; 9(5):2402–2413.

9. Zhou Z, Wang Y, Sun S, Zhang K, Wang L, Zhao H, et al. Paeonia lactiflora Pall. Polysaccharide alleviates depression in CUMS mice by inhibiting the NLRP3/ASC/Caspase-1 signaling pathway and affecting the composition of their intestinal flora. J Ethnopharmacol. 2023; 316:116716.

10. Jiang H, Li J, Wang L, Wang S, Nie X, Chen Y, et al. Total glucosides of paeony: A review of its phytochemistry, role in autoimmune diseases, and mechanisms of action. J Ethnopharmacol. 2020; 258:112913.

11. Yan BF, Chen X, Chen YF, Liu SJ, Xu CX, Chen L, et al. Aqueous extract of Paeoniae Radix Alba (Paeonia lactiflora Pall.) ameliorates DSS-induced colitis in mice by tunning the intestinal physical barrier, immune responses, and microbiota. J Ethnopharmacol. 2022; 294:115365.

12. Haworth M, Marino G, Atzori G, Fabbri A, Daccache A, Killi D, et al. Plant Physiological Analysis to Overcome Limitations to Plant Phenotyping. Plants (Basel). 2023; 12(23):4015.

13. Zeng ZH, Zhong L, Sun HY, Wu ZK, Wang X, Wang H, et al. Parallel evolution of morphological and genomic selfing syndromes accompany the breakdown of heterostyly. New Phytol. 2024; 242(1):302–316.

14. Xu S, Liu W, Liu X, Qin C, He L, Wang P, et al. DUS evaluation of nine intersubgeneric hybrids of *Paeonia lactiflora* and fingerprint analysis of the chemical components in the roots. Front Chem. 2023; 11:1158727.

15. Pace BA, Perales HR, Gonzalez-Maldonado N, Mercer KL. Physiological traits contribute to growth and adaptation of Mexican maize landraces. PLoS One. 2024; 19(2):e0290815.

16. Sun W, Yuan X, Liu ZJ, Lan S, Tsai WC, Zou SQ. Multivariate analysis reveals phenotypic diversity of Euscaphis japonica population. PLoS One. 2019; 14(7):e0219046.

17. Madhuvanthi CK, Eswaran M, Karthick T, Balasubramanian A, Dasgupta MG. Assessment of seed-and seedling-related traits in *Santalum album* (Indian sandalwood) reveals high adaptive potential. J Biosci. 2024; 49:13.

18. Saieed MAU, Zhao Y, Chen K, Rahman S, Zhang J, Islam S, et al. Phenotypic Plasticity of Yield and Yield-Related Traits Contributing to the Wheat Yield in a Doubled Haploid Population. Plants (Basel). 2023; 13(1):17.

19. Alía R, Climent J, Santos-Del-Blanco L, Gonzalez-Arrojo A, Feito I, Grivet D, et al. Adaptive potential of maritime pine under contrasting environments. BMC Plant Biol. 2024; 24(1):37.

20. Rahman MW, Deokar AA, Lindsay D, Tar’an B. Novel Alleles from *Cicer reticulatum* L. for Genetic Improvement of Cultivated Chickpeas Identified through Genome Wide Association Analysis. Int J Mol Sci. 2024; 25(1):648.

21. Komatsu K. Comprehensive study on genetic and chemical diversity of Asian medicinal plants, aimed at sustainable use and standardization of traditional crude drugs. J Nat Med. 2024; 78(2):267–284.

22. Chen Q, Chen L, Teixeira da Silva JA, Yu X. The plastome reveals new insights into the evolutionary and domestication history of peonies in East Asia. BMC Plant Biol. 2023; 23(1):243.

23. Abdullaeva Y, Ratering S, Rosado-Porto D, Ambika Manirajan B, Glatt A, Schnell S, et al. Domestication caused taxonomical and functional shifts in the wheat rhizosphere microbiota, and weakened the natural bacterial biocontrol against fungal pathogens. Microbiol Res. 2024; 281:127601.

24. Peng LP, Cai CF, Zhong Y, Xu XX, Xian HL, Cheng FY, et al. Genetic analyses reveal independent domestication origins of the emerging oil crop Paeonia ostii, a tree peony with a long-term cultivation history. Sci Rep. 2017; 7(1):5340.

25. Cetiz MV, Turumtay EA, Burnaz NA, Özhatay FN, Kaya E, Memon A, et al. Phylogenetic analysis based on the ITS, matK and rbcL DNA barcodes and comparison of chemical contents of twelve Paeonia taxa in Türkiye. Mol Biol Rep. 2023; 50(6):5195–5208.

26. Fan M, Li X, Zhang Y, Wu S, Song Z, Yin H, et al. Floral organ transcriptome in Camellia sasanqua provided insight into stamen petaloid. BMC Plant Biol. 2022; 22(1):474.

27. Cui L, Chen T, Zhao X, Wang S, Ren X, Xue J, et al. Karyotype Analysis, Genomic and Fluorescence In Situ Hybridization (GISH and FISH) Reveal the Ploidy and Parental Origin of Chromosomes in *Paeonia* Itoh Hybrids. Int J Mol Sci. 2022; 23(19):11406.

28. Yang Y, Sun M, Li S, Chen Q, Teixeira da Silva JA, Wang A, et al. Germplasm resources and genetic breeding of *Paeonia*: a systematic review. Hortic Res. 2020; 7:107.

29. Zhang J, Zhang D, Wei J, Shi X, Ding H, Qiu S, et al. Annual growth cycle observation, hybridization and forcing culture for improving the ornamental application of Paeonia lactiflora Pall. in the low-latitude regions. PLoS One. 2019; 14(6):e0218164.

30. Yang Z, Cao Y, Shi Y, Qin F, Jiang C, Yang S. Genetic and molecular exploration of maize environmental stress resilience: Toward sustainable agriculture. Mol Plant. 2023; 16(10):1496–1517.

31. Williams BR, Miller AJ, Edwards CE. How do threatened plant species with low genetic diversity respond to environmental stress? Insights from comparative conservation epigenomics and phenotypic plasticity. Mol Ecol Resour. 2023.

32. Farinon B, Picarella ME, Siligato F, Rea R, Taviani P, Mazzucato A. Phenotypic and Genotypic Diversity of the Tomato Germplasm From the Lazio Region in Central Italy, With a Focus on Landrace Distinctiveness. Front Plant Sci. 2022; 13:931233.

33. Guo Y, Jin GZ, Liu ZL. Effects of phyllotaxy on variation and inner relationships of leaflet traits in compound-leaved plants. Ying Yong Sheng Tai Xue Bao. 2023; 34(3):577–587.

34. Zhao W, Bai Q, Zhao B, Wu Q, Wang C, Liu Y, et al. The geometry of the compound leaf plays a significant role in the leaf movement of Medicago truncatula modulated by mtdwarf4a. New Phytol. 2021; 230(2):475–484.

35. Ouiyangkul P, Tantishaiyakul V, Hirun N. Exploring potential coformers for oxyresveratrol using principal component analysis. Int J Pharm. 2020; 587:119630.

36. Wang Q, Li X, Chen H, Wang F, Li Z, Zuo J, et al. Mapping combined with principal component analysis identifies excellent lines with increased rice quality. Sci Rep. 2022; 12(1):5969.

37. Zhou X, Wang D, Mao Y, Zhou Y, Zhao L, Zhang C, et al. The Organ Size and Morphological Change During the Domestication Process of Soybean. Front Plant Sci. 2022; 13:913238.

38. Guo L, Wang Y, da Silva JAT, Fan Y, Yu X. Transcriptome and chemical analysis reveal putative genes involved in flower color change in Paeonia ’Coral Sunset’. Plant Physiol Biochem. 2019; 138:130–139.

39. Zhou C, Wu H, Sheng Q, Cao F, Zhu Z. Study on the Phenotypic Diversity of 33 Ornamental *Xanthoceras sorbifolium* Cultivars. Plants (Basel). 2023; 12(13):2448.

40. Shi S, Wang E, Li C, Zhou H, Cai M, Cao C, et al. Comprehensive Evaluation of 17 Qualities of 84 Types of Rice Based on Principal Component Analysis. Foods. 2021; 10(11):2883.

